# SSP: An R package to estimate sampling effort in studies of ecological communities

**DOI:** 10.1101/2020.03.19.996991

**Authors:** Edlin J. Guerra-Castro, Juan Carlos Cajas, Nuno Simões, Juan J Cruz-Motta, Maite Mascaró

## Abstract

`SSP` (simulation-based sampling protocol) is an R package that uses simulation of ecological data and dissimilarity-based multivariate standard error (*MultSE*) as an estimator of precision to evaluate the adequacy of different sampling efforts for studies that will test hypothesis using permutational multivariate analysis of variance. The procedure consists in simulating several extensive data matrixes that mimic some of the relevant ecological features of the community of interest using a pilot data set. For each simulated data, several sampling efforts are repeatedly executed and *MultSE* calculated. The mean value, 0.025 and 0.975 quantiles of *MultSE* for each sampling effort across all simulated data are then estimated and standardized regarding the lowest sampling effort. The optimal sampling effort is identified as that in which the increase in sampling effort do not improve the precision beyond a threshold value (e.g. 2.5 %). The performance of `SSP` was validated using real data, and in all examples the simulated data mimicked well the real data, allowing to evaluate the relationship *MultSE* – *n* beyond the sampling size of the pilot studies. `SSP` can be used to estimate sample size in a wide range of situations, ranging from simple (e.g. single site) to more complex (e.g. several sites for different habitats) experimental designs. The latter constitutes an important advantage, since it offers new possibilities for complex sampling designs, as it has been advised for multi-scale studies in ecology.

## Background

Defining sample size is a key decision in the planning of ecological research. In the context of hypothesis testing, a decision to take too few samples could produce misleading information about the statistical population, imprecise statistics or a high probability of retaining a false null hypothesis. Instead, increasing sampling size improves the precision of estimations and the power of statistical tests, but will also increase its costs (Mapstone 1995, Underwood 1997, Underwood and Chapman 2003). There is a wide variety of methodologies aimed at optimizing the use of resources to obtain the best possible sampling design at the lowest cost (a.k.a cost-benefit optimization). Most of the methodologies used for this purpose, however, are based on statistical theories that takes into account only one response variable; that is, when the variable of interest can be represented as a single descriptor. In early ecological studies at community level, sample sizes were usually determined on the basis of single merging variables such as species richness or some sort of diversity index (Green 1979, Clarke and Green 1988). However, the study of communities has evolved from the use of univariate descriptors to the use of multivariate/dissimilarity-based statistics (Clarke 1993, Anderson et al. 2006, Legendre and De Cáceres 2013). Therefore, the analytical methods for estimation of sample size should consider the highly-dimensional structure of ecological data (Anderson and Santana-Garcon 2015, Blanchet et al. 2016).

Most of the conventional approaches in experimental and sampling design consist on evaluating the precision of the arithmetic mean of a response variable obtained in a previous pilot study in order to estimate how many replicas are needed to improve that precision. These strategies consider the expected random error, a previous estimate of the natural variability of the variable, the effort with which such information was obtained and some theoretical distribution as a reference (e.g. standardized normal distribution) (Quinn and Keough 2002). Statistical precision refers to the level of concordance between multiple estimators of the same parameter under the same sampling procedure (Underwood 1997). To estimate the precision of a mean obtained from a random sample, the standard error of the mean must be calculated 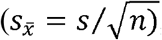, where *s* represents the standard deviation of the sample and *n* the number of sampling units. Data from a previous pilot study providing such information is often required. Because the denominator of the standard error is the sample size, the standard error will always decrease (and precision will improve) as the sampling effort increases.

Anderson and Santana-Garcon (2015) developed a computationally intensive statistical approach that allows the estimation of a multivariate *pseudo* standard error (*MultSE*), as a proxy of precision, in order to identify an optimal sample size (eqn (3) Anderson and Santana-Garcon 2015). This approach consists of a double resampling (with and without replacement) of a data matrix obtained during a pilot study. For each resampling, the dissimilarities between each pair of samples of a randomly chosen subset of the main matrix are estimated and the *MultSE* is calculated. The procedure is repeated several times with *n_i_* = 2, 3, 4 to *n*, where *n* refers to the original sample size in the pilot study. The behaviour of *MultSE* is projected on a plot of means and error bars, with the abscissa showing the sampling effort and the ordinate showing *MultSE* values (see Figure 1 in Anderson and Santana-Garcon 2015). From the plot, the way precision relates to effort can be observed, thereby helping to identify the optimum sampling effort for an acceptable measure of standard error. The R scripts for this approach are available as supplementary material in Santana-Garcon (2015).

**Figure 1.**
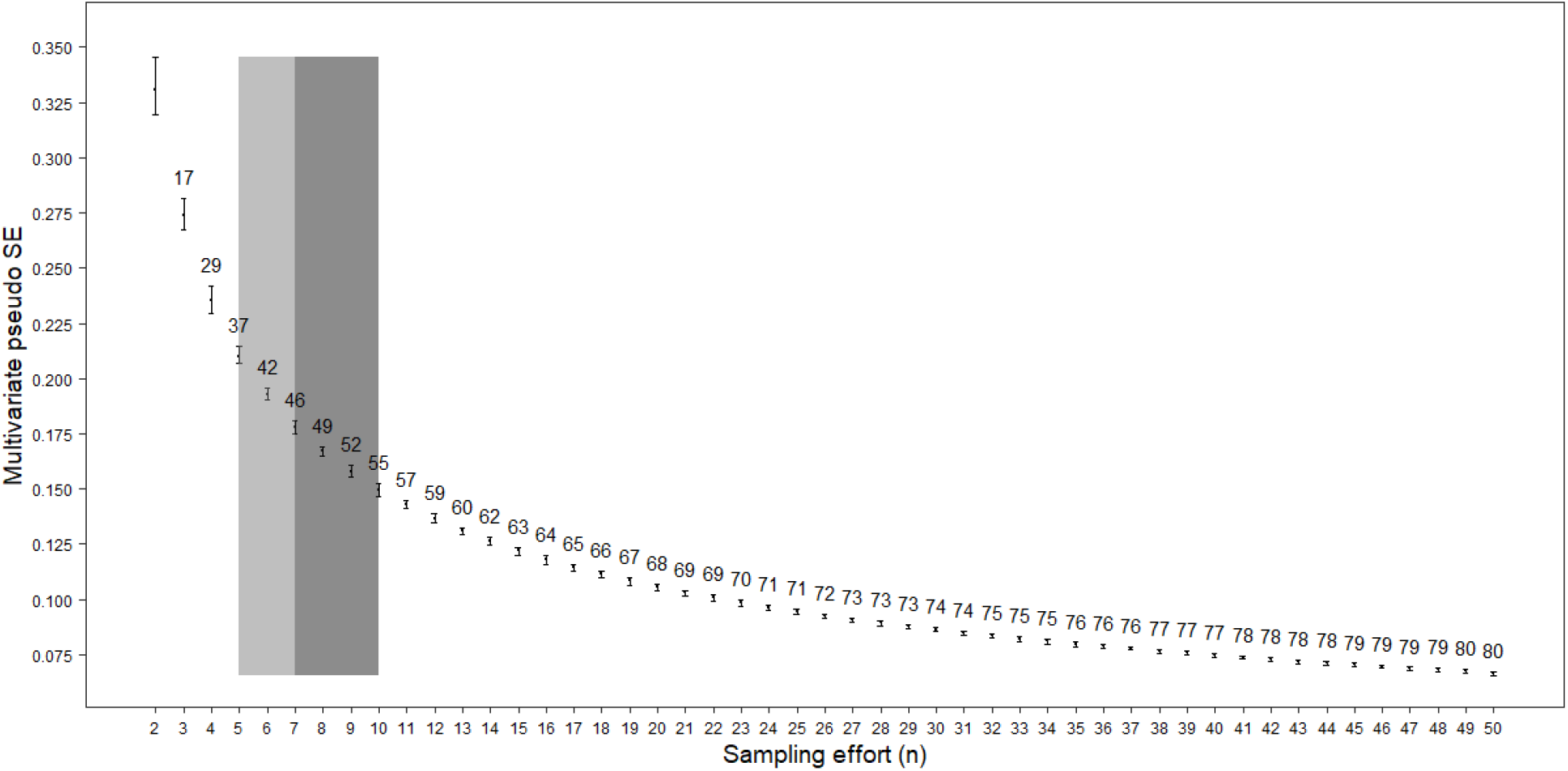
Example 1: *MultSE* and sampling effort relationship using micromollusk simulated data. The shaded areas indicate the range of samples in which each increase in sampling effort provides suboptimal improvement (light gray), and optimal improvement (dark gray).

We present `SSP`, an R package (R Core Team, 2013) designed to estimate sample effort in studies of ecological communities based on the definition of *MultSE* (Anderson and Santana-Garcon 2015), but sampling intensively over several sets of simulated data instead of the double resampling approach over a unique pilot data set. In general, the protocol consists of 1) simulating several data matrixes that retain observed properties of the community of interest, 2) obtaining independent estimates of *MultSE* from those simulated data matrixes, for different sample sizes and number of sites, and 3) a quantitative identification of the optimal sampling effort as well as the graphic representation of the *MultSE – n* relationship and the optimal effort. Data collected in a standard pilot survey are used to simulate the data matrixes but are not included in the resampling procedure. With `SSP`, users will be able to objectively identify the number of samples and sites necessary to characterize the community of interest with sufficient precision at a reasonable cost. The use of several simulated larger data matrixes as a central element of the procedure will assure a better appreciation of the *MultSE – n* relationship over a wider range of sample sizes. `SSP` was evaluated using three sets of real data: (1) micromollusk of marine shallow sandy bottoms (2) coral reef sponges; and (3) epibenthic assemblages on Caribbean mangrove roots.

## Methods and features

### Workflow of SSP

`SSP` was divided in seven stages: (*i)* extrapolation of assemblage parameters using pilot data, (*ii*) simulation of several data sets based on extrapolated parameters, (*iii*) evaluation of plausibility of simulated data, (*iv*) repeated estimations of *MultSE* for different sampling designs in simulated data sets, (*v*) summary of the behaviour of *MultSE* for each sampling design across all simulated data sets, (*vi*) identification of the optimal sampling effort, and (*vii*) graphical representation of the *MultSE* and the sampling effort. For each of these steps, we provide seven functions, these are:

i. `assempar`: The following ecological properties of the assemblage are estimated: potential number of species, probability of occurrence of each species within and among sites, the patterns of abundances of each species, and the patterns of spatial aggregation of species. The potential number of species in the assemblage is estimated with any of the incidence-based nonparametric methods available in the specpool function of the vegan package (Oksanen et al. 2015). The probability of occurrence of each species is calculated between and within sites. The former is computed as the frequency of occurrences of each of the species against the number of sites sampled, the second as the weighted average frequencies in sites where the species were present (Gaston 1994, Magurran and Henderson 2011). The mean (and variance) of the abundance of each species was also estimated to be modelled with the appropriate statistical distribution (McArdle and Anderson 2004). The degree of spatial aggregation of species (only for real counts of individuals), was identified with the index of dispersion *D* (Clarke et al. 2006). The corresponding properties of unseen species were approximated using information of observed species. Specifically, the probabilities of occurrence were assumed to be equal to the rarest species of pilot data. The mean (and variance) of the abundances are defined using random Poisson values with lambda as the overall mean of species abundances. `assempar` returns an object of class list, to be used by `simdata`.
ii. `simdata`: The simulation starts by setting the dimensions of data matrix *Ŷ* (i.e. number of columns and rows). The number of columns was programmed to be equal to the potential number of species, while the number of rows (*N_t_*) is defined arbitrarily as the potential number of sampling units per site (*N*) multiplied by the potential number of sites (*M*). The presence/absence of each species at each site are simulated with Bernoulli trials and probability of success equals to the empirical frequency of occurrence of each species among sites in the pilot data. Then, for sites with the simulated presence of the species, Bernoulli trials are used again with probability of success equal to the empirical frequency estimated within the sites in pilot data. If required, the presence/absence matrices are converted to matrices of abundances replacing presences by random values from an adequate statistical distribution and parameters equals to those estimated in the pilot data. Counts of individuals are generated using Poisson or negative binomial distributions, depending on the degree of aggregation of each species in the pilot data (McArdle and Anderson 2004, Anderson and Walsh 2013). When abundances were measured as a continuous variable (i.e. coverage, biomass), they are generated using the lognormal distribution. The simulation procedure is repeated to generate as many simulated data matrices as needed. It is important to highlight that the procedure assumes ‘homogeneous’ environmental conditions across samples; this is, that multivariate structure of the assemblage is produced by intrinsic properties of species (e.g. patterns of gregariousness/dispersion) and not by environmental constrains. This condition means that simulations should not be done combining data from different habitats (e.g. mixing quadrats from hide and low tides in a rocky shore or along an environmental gradient). For these cases, simulations should be done independently for each habitat. This function returns an object of class list that will be used by `sampsd` and `datquality`.
iii. `datquality`: The quality of the simulated data matrixes is assessed by the resemblance these had to the pilot data considering the following estimations: (i) the average number of species per sampling unit, (ii) the average species diversity (Simpson diversity index) per sampling unit, and (iii) the multivariate dispersion (MVD), measured as the average dissimilarity from all sampling units to the main centroid in the space of the dissimilarity measure used (Anderson 2006). In general, it is desirable that (i), (ii) and (ii) be similar between simulated and pilot data.
iv. `sampsd`: If several virtual sites have been simulated, subsets of sites of size 2 to *m* are sampled, followed by the selection of sampling units (from 2 to *n*) using inclusion probabilities and self-weighted two-stage sampling (Tillé 2011). Each combination of sampling effort (number of sample units and sites), are repeated several times (e.g. 100) for all simulated matrixes. If simulated data correspond to a single site, sampling without replacement is performed several times (e.g. 100) for each sample size (from 2 to *n*) within each simulated matrix. This approach is computationally intensive, especially when *k* is high. Keep this in mind because it will affect the time to get results. For each sample, suitable pre-treatments are applied, and distance/similarity matrixes estimated using the appropriate coefficient. When simulations were done for a single site, the *MultSE* is estimated as 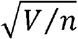, being *V* the *pseudo* variance measured at each sample of size *n*. When several sites were generated, *MultSE* are estimated using the residual mean squares 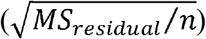 and the sites mean squares 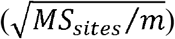 from a distance-based multivariate analysis of variance.
v. `summary_ssp`: This function is required to estimate an average of all *MultSE* obtained with the *k* repetitions for each sampling effort within each simulated data, and then an overall mean as well as the lower and upper intervals of means for each sampling effort among all simulated data. In order to have a general and comparable criteria to evaluate the rate of change of the averaged *MultSE* with respect to the sampling effort, a relativization to the maximum *MultSE* value (obtained with the lower sampling effort) is calculated; then, a standard forward finite derivation is computed. All these results are presented in a data frame that will be used to plot *MultSE* with respect to the sampling effort, using `ssp_plot`.
vi. `ioptimum`: This function identifies three cut-off points based on the finite derivative between the standardized *MultSE* and the sampling effort (as the percentage of improvement in precision per sample unit, by default 10%, 5%, and 2.5%), allowing us to identify: (1) the minimum improvement required, (2) sub-optimal improvement, and (3) optimal improvement. It is possible that the cut-off points defined by default will not be achieved, which might occur if the arguments *n* or *m* of `sampsd` were set small. In situations like this, a warning message will specifically indicate which cut-off point was not achieved and will return the maximum effort currently executed. To achieve these improvements, `sampsd` and `summary_ssp` must be run again with higher values of *n* or *m*. Alternatively, the cut-off points can be made flexible, say, 15%, 10%, 5%, respectively, or higher.
vii. `ssp_plot`: This function allows to visualize the behavior of the *MultSE* as sampling effort increases. When the simulation involves two sampling scales, a graph for samples and one for sites will be generated. Above the *MultSE*~Sampling effort projection, two shaded areas are drawn, highlighting: sub-optimal improvement (light gray), and optimal improvement (dark gray). Both reflect the sampling effort that improves the precision at acceptable (light gray) or desirable levels (dark gray), but beyond the later, any gain could be considered unnecessary. In addition, for each sampling effort, the relativized improvement (in relation to the *MultSE* estimated with the lower sampling effort) is presented cumulatively (as percentages). This is very useful because it indicates exactly how much the precision is improved for each increase in sampling units. The graphic is generated with ggplot2, so the resulting object can be modified using functions of that package

### Examples

1. Micromollusks of marine shallow sandy bottoms: Presence/absence of 68 species were registered in six cores of 10 cm diameter and 10 cm deep taken in sandy bottoms around Cayo Nuevo, Gulf of Mexico, Mexico (a small reef cay located 240 km off the North-Western coast of Yucatan). Data correspond to a study on the biodiversity of marine benthic reef habitats off the Yucatan shelf (Ortigosa et al. 2018). The main objective was to estimate an adequate sampling effort for further quantitative studies to characterize the temporal changes in species composition. The R script for this example is in box Example 1. To speed up the process, only 20 data sets were simulated, each data matrix consisted of *N* = 100 potential sampling replicates in one site, and sample size’s subsets from 2 to 50 were repeated 10 times. The Jaccard index was used as the similarity measure between sample units. `SSP` indicates that with a sampling effort between 5 and 10 samples, the worst precision would be improved from 37% (suboptimal) to 55% (optimal), with remarkable precision gained with each additional sample (Fig. 1). After 11 samples, the improvement obtained with the increment of the sampling effort is small enough to consider the extra effort unnecessary and redundant.

**Example 1.**
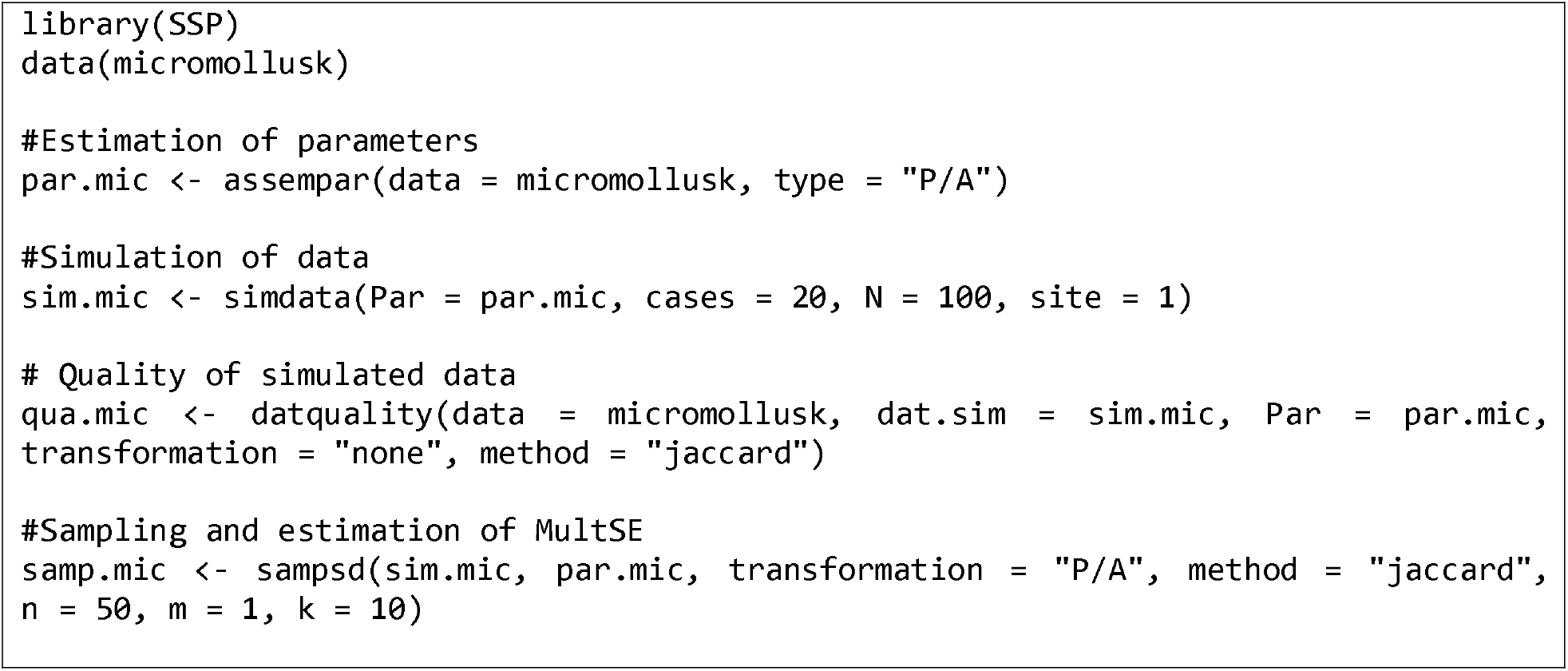
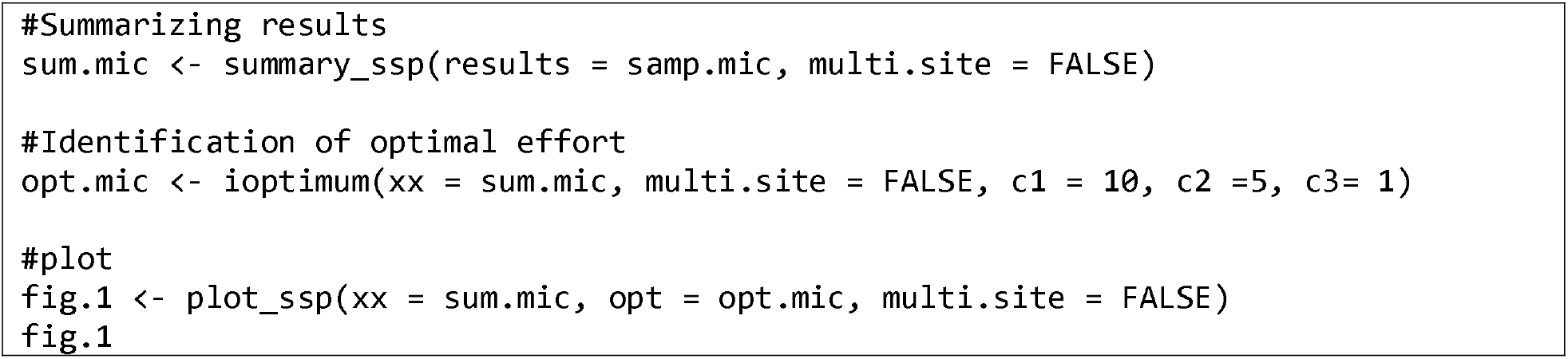
Aplication of `SSP` to micromollusks assemblage from sandy bottoms around Cayo Nuevo, Gulf of Mexico, Mexico (Ortigosa et al. 2018)

2. Coral reef sponges: Structure and composition of sponge assemblages associated with reefs from Alacranes Reef National Park (ARNP), Gulf of Mexico, Mexico, was estimated in 36 transects of 20 m × 1 m across 6 sites (≈ 4 to 5 transect per site). In each transect, the colonies of 41 species of sponges were counted. This data corresponds to a pilot study on sponge biodiversity in reef habitats of the Yucatán shelf (Ugalde et al. 2015). The main objective was to estimate an adequate sampling effort at two spatial scales (i.e. transect and sites) for further quantitative studies. The studied area represented the leeward area of the reef, with very similar reef geomorphology. Therefore, it cannot be argued a priori that spatial differences in sponge diversity (if any) were due to some environmental model. In this sense, we considered valid to simulate data for the entire leeward area, using the information of the six sites. Again, to speed up the process, only 10 data sets were simulated, each consisting in 20 virtual sites with 20 virtual transects per site. Each combination of *n* (from 2 to 20) and sites (from 2 to 20) were repeatedly sampled 10 times. The Bray-Curtis index was used as the similarity measure between sample units after square root transformation of simulated abundances. The R script for this second example is in box 2.

**Example 2.**
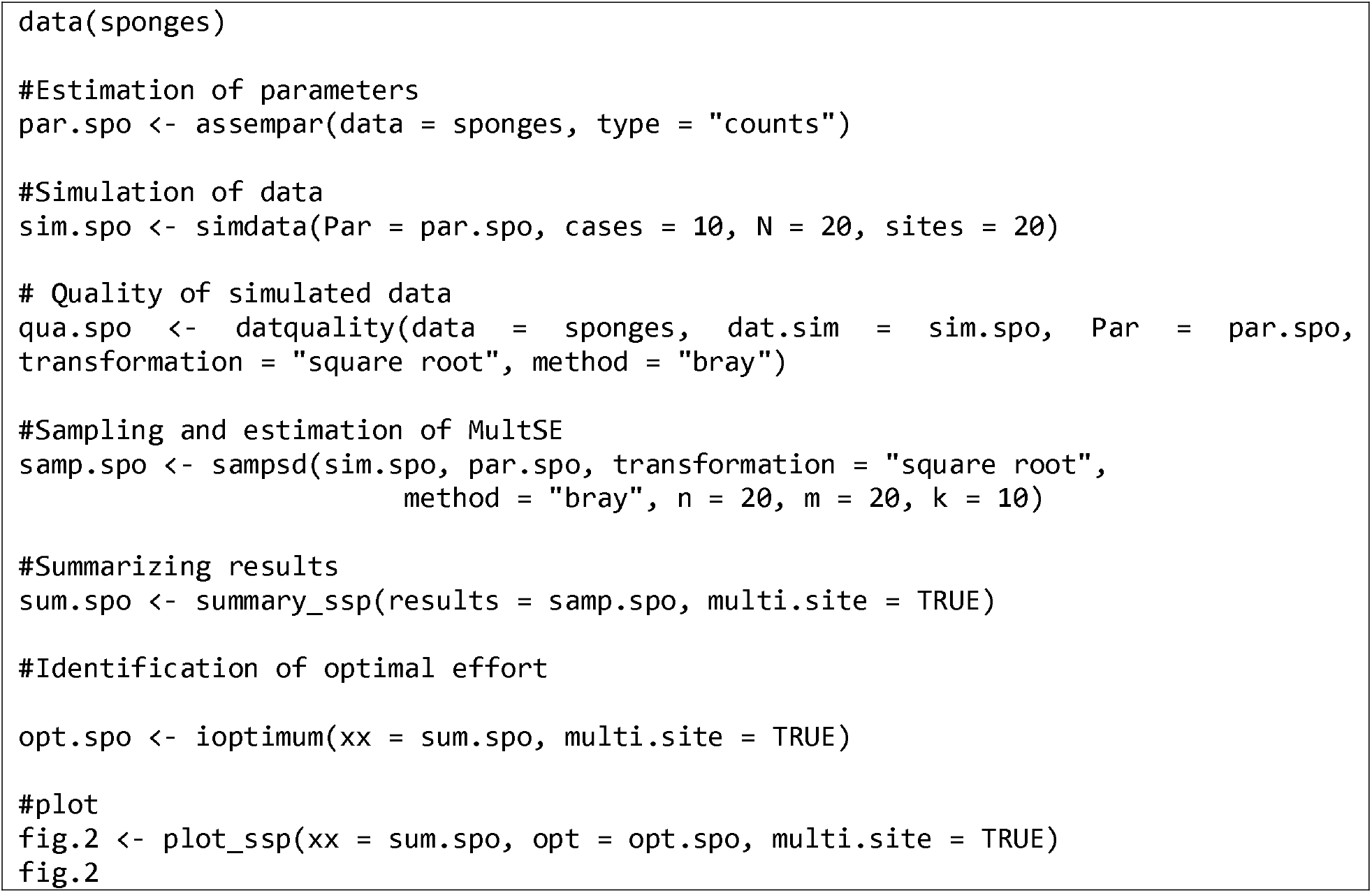
Aplication of `SSP` to sponge assemblages associated with reefs from Alacranes Reef National Park (ARNP), Gulf of Mexico, Mexico (data from Ugalde et al. 2015)

At the scale of sites, there was a noticeable decrease of the *MultSE* between 5 and 11 sites (Fig. 2). This range of sampling is represented by two shaded areas: light gray) and optimal improvement (dark gray), respectively. The first represent improvements of 44% with 7 sites, the second 55% with 11 sites. Sampling 9 sites will improve the worst precision in approximately 51%. At the scale of transect, the suboptimal improvement is accomplished with 5 or 6 replicates, whereas the optimal is beyond 7 replicates, but below 11. It can also be noted that each additional sample improves the highest *MultSE* by 2-3%, achieving a 55% improvement with 10 transects. It should be pointed out the marked differences in *MultSE* obtained for the two sources of variation. The magnitude of the difference in variation fits that obtained in the *pseudo*- components of variation estimated for sites and residuals in a distance-based multivariate analysis of variance of the pilot data σ_sites_ = 26.4, σ_transects_ = 34.7 (R script in the Supplementary material Appendix 1). Based on these results, and considering the costs (time and/or money) of visiting a site and doing each transect, as well as the relative importance of each spatial scale, it will be convenient to keep the number of sites within suboptimal improvement (i.e. 5-7), and set the number of transects to 8.

**Figure 2.**
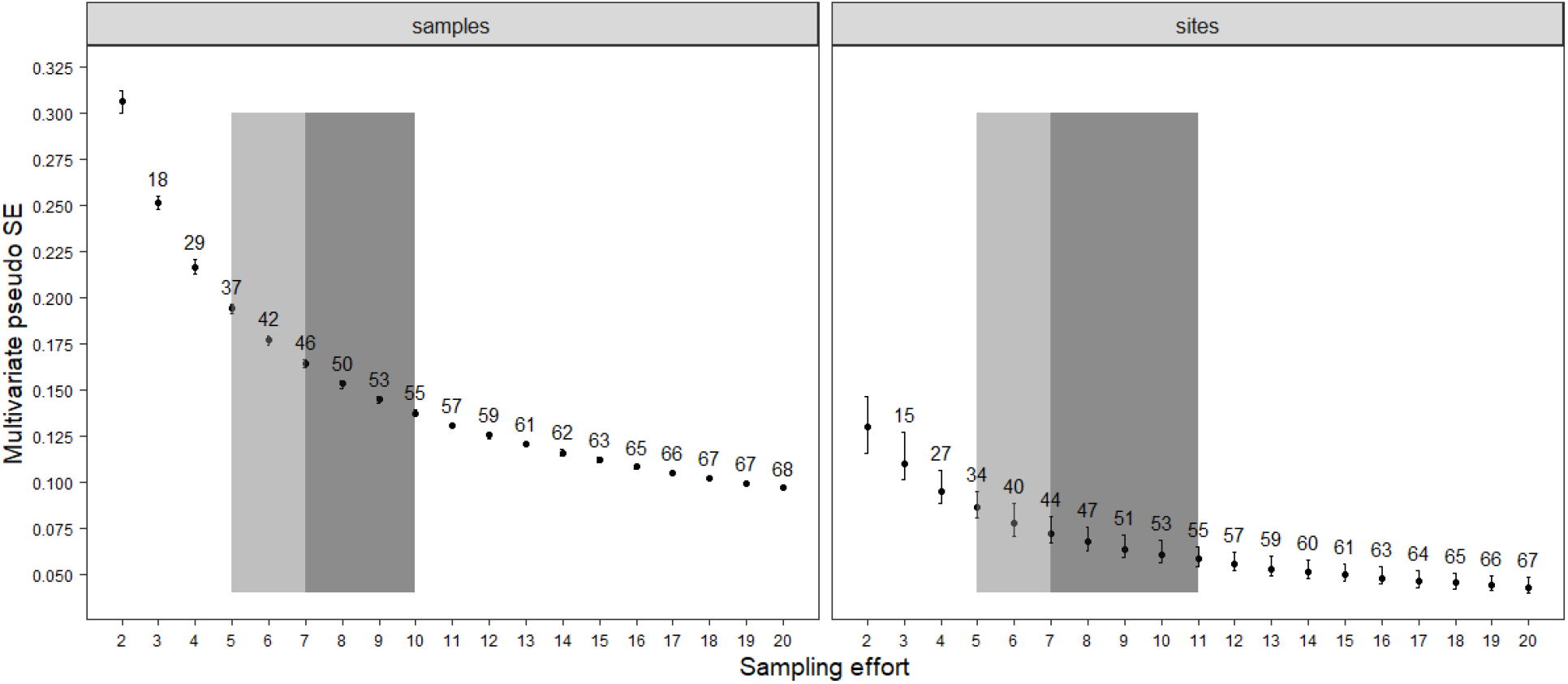
Example 2: *MultSE* and sampling effort relationship using sponge simulated data. The shaded areas indicate the range of samples in which each increase in sampling effort provides suboptimal improvement (light gray), and optimal improvement (dark gray).

3. Epibenthic assemblages on Caribbean mangrove roots: Data consists of the coverage (by point-intercept) of 116 taxa identified in 180 mangrove roots, sampled under a hierarchically nested spatial design that included six random sites within each of three sectors of the lagoon system corresponding to a strong environmental gradient: external (E), intermediate (M), and internal (I). The abundance of epibenthic organisms of 10 roots were described within each site, producing a total of 60 roots in each sector. One of the main objectives of this pilot study was to define a sampling effort to evaluate spatiotemporal patterns of variation in species composition among sectors considering the environmental gradient. To achieve this, it was necessary to identify a sampling effort (number of sites and roots) that guarantee the best precision at the lowest cost. For each sector, 20 data sets were simulated, each with 30 virtual sites and 30 virtual roots each. Then, two-stage random sampling with sites from 2 to 20 and roots from 2 to 20 were repeated 10 times for each combination. The Bray-Curtis index was used as the similarity measure between sample units after fourth root transformation of abundances. The R script for all operations is in the box Example 3.

**Example 3.**
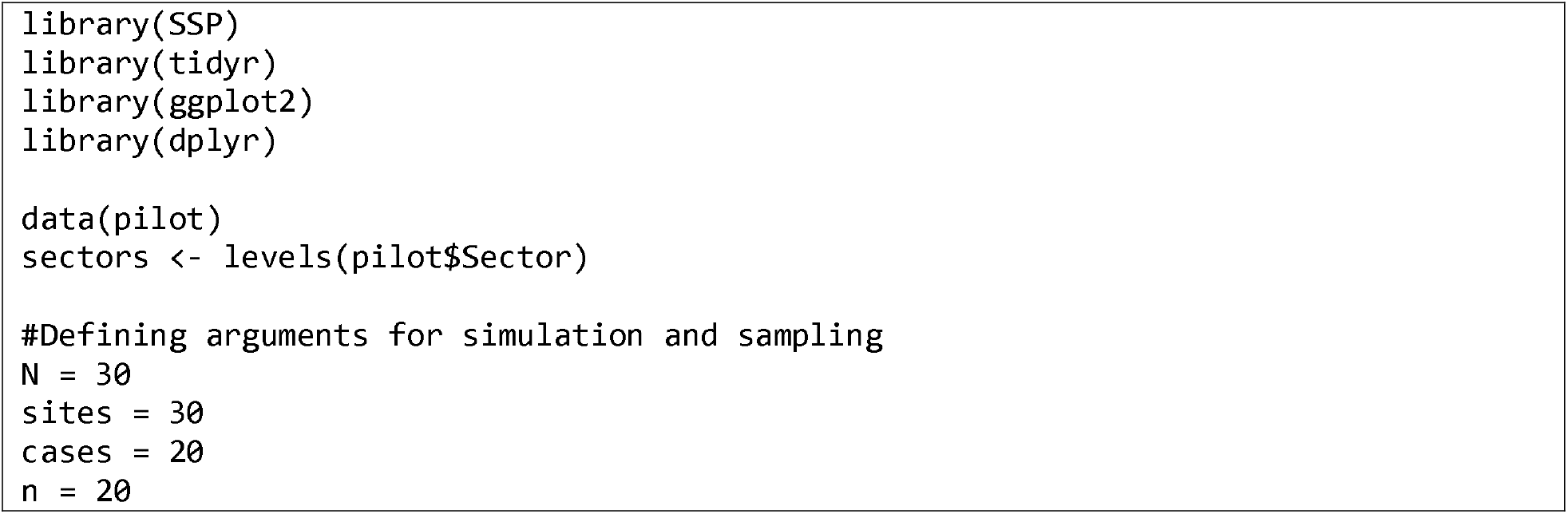
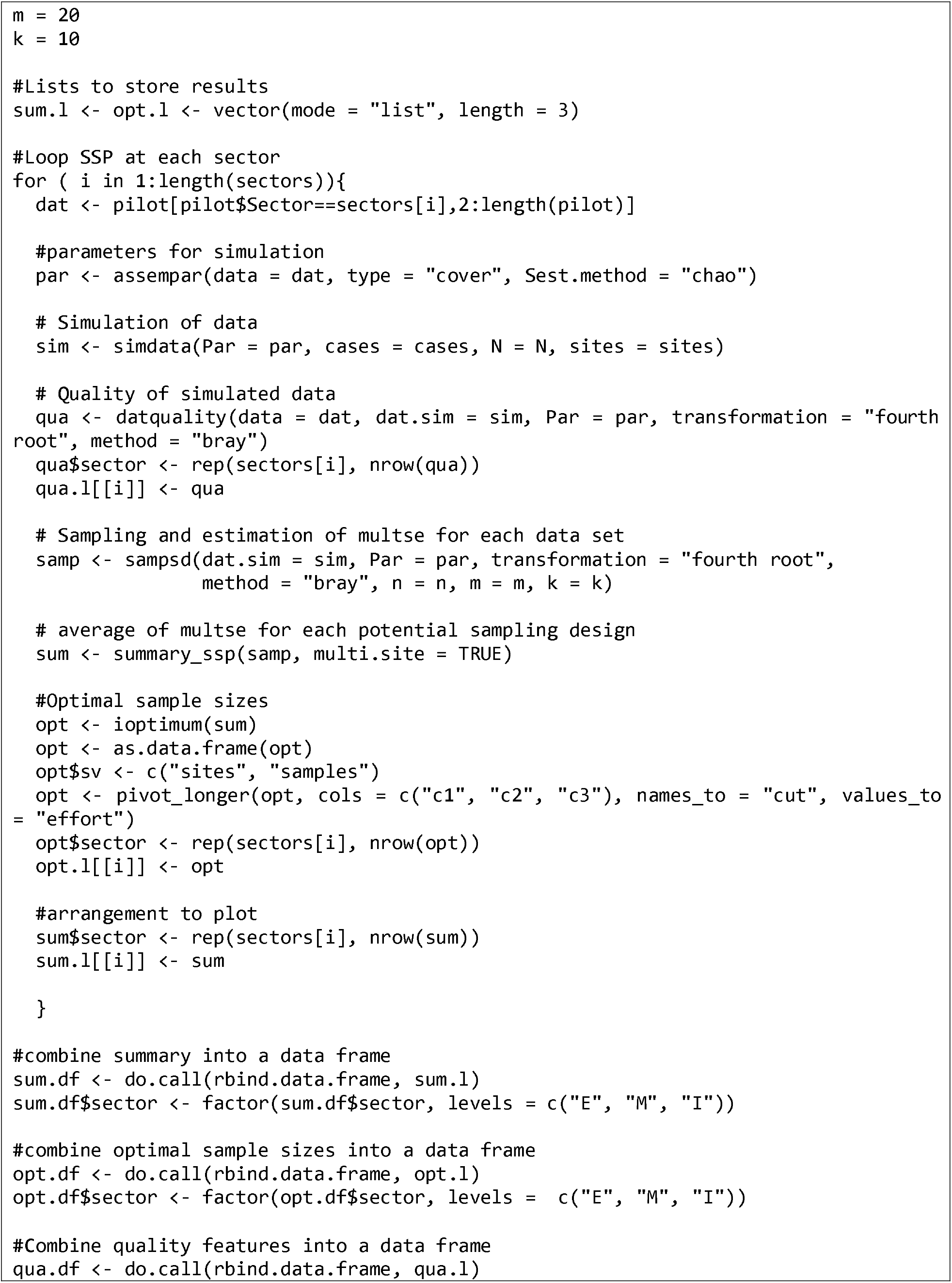
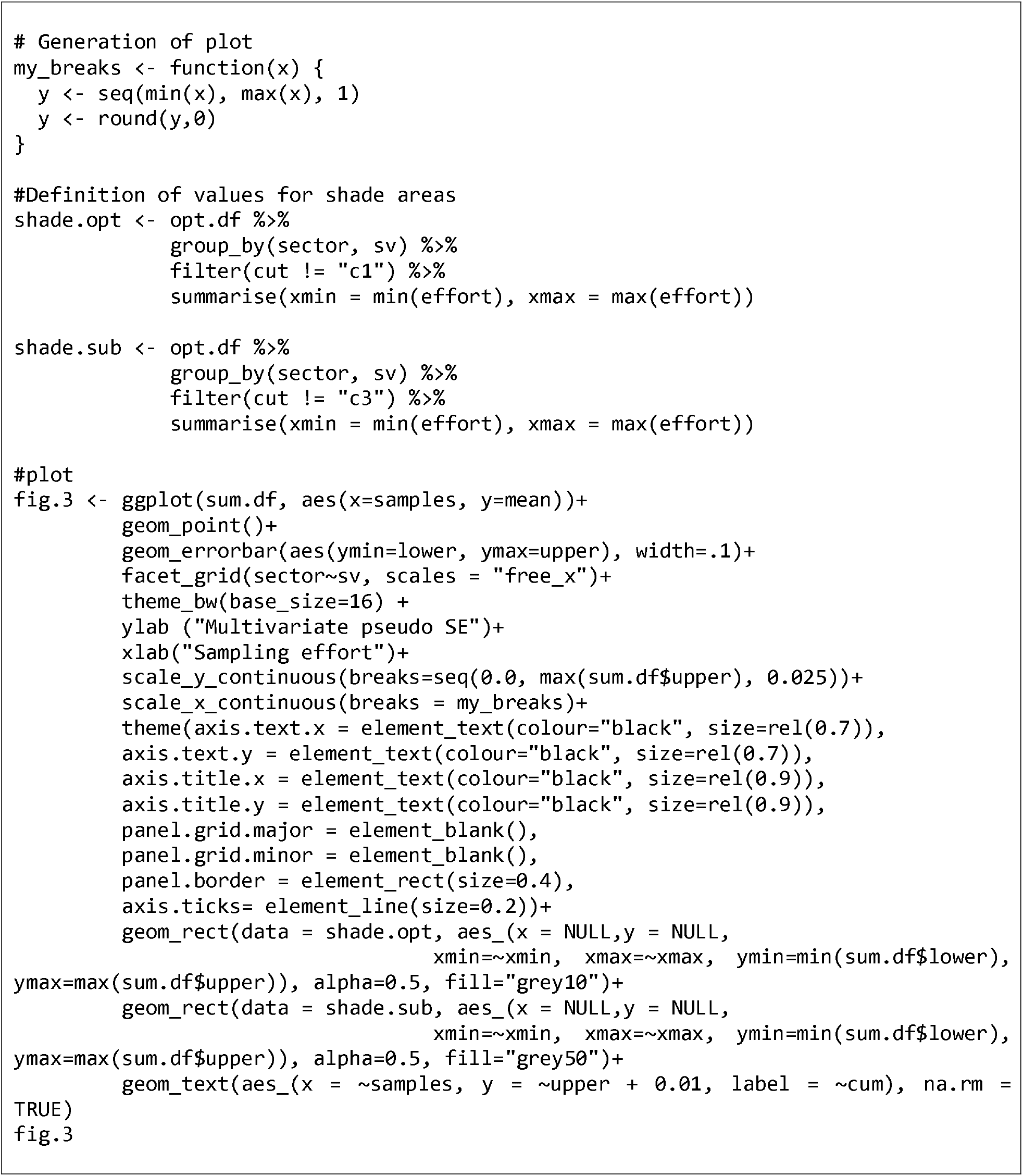
Aplication of `SSP` to epibenthic assemblages on Caribbean mangrove roots (data from Guerra-Castro et al., 2011). Considering the environmental differences between each sector, `SSP` was applied independently to each sector. The results were merged and analysed simultaneously.

`SSP` indicate that the number of sites required to obtain the same precision is similar among sectors, in all cases the *MultSE* was below 0.1. Considering the *MultSE* values estimated in other studies (Anderson and Santana-Garcon 2015), 0.1 is a good level of precision even for a few sites, making suboptimal improvement a good criterion for defining sampling effort for this spatial scale. On the other hand, the variability among roots was considerable higher in all sites, between 0.245 and 0.287; consequently, to reduce the *MultSE* in half it is required an optimal improvement, at least 8 roots per site (Fig. 3). Indeed, in the original study, using a very primitive version of `SSP`, the sampling effort was defined with 4 sites per sector and 8 roots per site. This sampling effort was used by Guerra-Castro et al. (2016), and it had the statistical power to detect differences along the gradient despite reduced sampling effort.

## Discussion

The R package described in the present study expands the method developed by Anderson and Santana-Garcon (2015), but maintains its purpose as a quantitative tool to define sample size for studies of communities with data to be analysed using distance-based methods. Results herein show that `SSP` improves the usefulness of the original procedure regarding the following aspects: (1) *MultSE* – *n* relationship could be evaluated beyond the original sampling size of the pilot study; (2) *MultSE* estimates are obtained by sampling over several simulated data sets, ensuring statistical independence of the estimations; (3) sampling effort can be defined in terms of sub-optimal and optimal improvement regarding the highest *MultSE*, (4) this protocol can be used to estimate sample size in an wide range of situations from simple (i.e. sampling a single or few sites) to more complex experimental designs (i.e. sampling several sites for different habitats). The latter constitutes an important advantage, since it offers new possibilities for the planning of complex sampling designs, as it has been advised for multi-scale studies in ecology (Underwood and Chapman 1998, Leibold et al. 2004, Chase et al. 2018).

Data sets used in this study comprised a variety of sampling designs, in all of which the `SSP` package provided an appropriate and useful visual tool to identify the optimal sampling effort. The `SSP` protocol applied on data from the simplest case in Micromollusks of Cayo Nuevo showed that the sampling effort should be increased compared to the pilot study (Fig. 2). The sponge case brings the opportunity to evaluate the precision required considering a more complex situation, since the `SSP` protocol can estimate *MultSE* in two different spatial scales: transects and sites. The possibility of targeting sampling effort in this way constitutes a key advantage in experimental design, since it allows a revaluation in the number of sampling sites, hence fieldwork costs. Similarly, results of the simulation on the data from epibenthic fauna in mangrove roots demonstrated the potential of `SSP` to define sampling effort in studies of communities in heterogeneous environments. Decisions such as these have a direct impact on the costs of a sampling project, and an adequate combination of effort at different spatial scales will help optimize the allocation of often limited resources. Optimization of sampling designs using cost-benefit procedures described by previous authors (e.g. Underwood 1997) could easily be combined with the protocol presented here, further improving cost-effective allocations of sampling effort.

It is important to mention that `SSP` is not free from assumptions. Simulations do not consider any environmental constraint, neither co-occurrence structure of species. It is assumed that potential differences in species composition/abundance among samples and sites are mainly due to spatial aggregation of species, as estimated from the pilot data. Hence, any ecological property of the assemblage that was not captured by the pilot data, will not be reflected in the simulated data. Associations among species can be modelled using copulas, as suggested by Anderson et al. (2019), which could be included in an upcoming version of `SSP`.

Finally, although the protocol performs well with small pilot data, it is recommendable that the pilot sampling is not restricted to a small number of samples and captures the greatest possible variability in the system under study. After evaluating the quality of the simulated data with `datquality`, it is notable that it was not always possible to simulate data with properties identical to the pilot data (Table 1). Specifically, in Example 1, the average number of species per sample was considerably higher than the original data, this may be a consequence of the limited probability of occurrence for each species restricted number of samples. In the other examples, the simulated data fit well with respect to the pilot data, and these properties were achievable because the pilot data were extensive. In conclusion, if the properties of the simulated data resemble the community of interest and show ecological plausibility, the extrapolations derived from `SSP` will hold valid to define the sampling size of any study based on dissimilarity-based multivariate analysis.

**Table 1.** Output of `datquality` for the three examples: relevant features of original and simulated data. Features include the number of sample units (n), the number of sites (m), average number species/sample (S̅), average Simpson Diversity Index (aSDI) and range of multivariate dispersion (MVD).

| | | n | m | $\bar{S}$ | aSDI | MVD |
| --- | --- | --- | --- | --- | --- | --- |
| Micromolusk from Cayo Nuevo, Gulf of Mexico |  |  |  |  |  |  |
| | Pilot | 6 | 1 | 25.3 ($\pm 7.2$) | -- | 0.25 |
| | Simulated | 1000 | 1 | 30.5($\pm 3.9$) | -- | 0.22 - 0.24 |
| Sponges from Alacranes Reef, Gulf of Mexico |  |  |  |  |  |  |
| | Pilot | 4 | 8 | 12.4($\pm 4.3$) | 0.84($\pm 0.08$) | 0.16 |
| | Simulated | 20 | 20 | 9.1($\pm 2.4$) | 0.72($\pm 0.13$) | 0.22 - 0.23 |
| Epibionts on Caribbean mangrove roots |  |  |  |  |  |  |
| External sector | Pilot | 60 | 6 | 18.3($\pm 5.7$) | 0.82($\pm 0.07$) | 0.19 |
| | Simulated | 30 | 30 | 18.4($\pm 3.3$) | 0.90($\pm 0.03$) | 0.18 - 0.19 |
| Intermediate sector | Pilot | 60 | 6 | 16.1($\pm 5.25$) | 0.80($\pm 0.10$) | 0.18 |
| | Simulated | 30 | 30 | 16.2($\pm 2.9$) | 0.93($\pm 0.01$) | 0.14 - 0.15 |
| Internal sector | Pilot | 60 | 6 | 14.3($\pm 2.6$) | 0.83($\pm 0.07$) | 0.14 |
| | Simulated | 30 | 30 | 14.8($\pm 2.7$) | 0.86($\pm 0.05$) | 0.14 - 0.15 |

### Software availability

`SSP` is free and open source, available on GitHub (<https://github.com/edlinguerra/SSP>). Data from all three examples are available in the package. To cite `SSP` or acknowledge its use, cite this software note including the appropriate version number.

## Acknowledgements

EGC was supported by a DGAPA Post-doctoral Fellowship at the Universidad Nacional Autónoma de México (UNAM). This research was financed by the research project PE207416 (PAPIME, UNAM) under the supervision of MM. Funding was also provided by project #325 (CÁTEDRAS, CONACYT), the HARTE Research Institute for Gulf of Mexico Studies and the Harte Charitable Foundation. NS holds the Furgason Fellowship International Chair for Coastal and Marine Studies in Mexico. Special thanks to M. J. Anderson and several anonymous reviewers for providing constructive critical comments that substantially improved an early version of this manuscript.

## Authors’ contributions

Idea was conceived by EJGC and MM. Modelling and data analyses were carried out by EJGC and JCC. NS and JJCM provided data sets from the Gulf of Mexico and Venezuela, respectively. EJGC and MM wrote the first draft of the paper and JCC, NS and JJCM contributed substantially to revisions.

