## Supplementary material for "SSP: An R package to estimate sampling effort in studies of ecological communities"

#Example 1
library(SSP)
data(micromollusk)

#Estimation of parameters
par.mic <- assempar(data = micromollusk, type = "P/A")

#Simulation of data
sim.mic <- simdata(Par = par.mic, cases = 20, N = 100, site = 1)

# Quality of simulated data
qua.mic <- datquality(data = micromollusk, dat.sim = sim.mic, Par = par.mic, transformation = "none", method = "jaccard")

#Sampling and estimation of MultSE
samp.mic <- sampsd(sim.mic, par.mic, transformation = "P/A", method = "jaccard", n = 50, m = 1, k = 10)

#Summarizing results
sum.mic <- summary_ssp(results = samp.mic, multi.site = FALSE)

#Identification of optimal effort
opt.mic <- ioptimum(xx = sum.mic, multi.site = FALSE)

#plot
fig.1 <- plot_ssp(xx = sum.mic, opt = opt.mic, multi.site = FALSE)
fig.1

#Example 2
data(sponges)

#Estimation of parameters
par.spo <- assempar(data = sponges, type = "counts")

#Simulation of data
sim.spo <- simdata(Par = par.spo, cases = 10, N = 20, sites = 20)

# Quality of simulated data
qua.spo <- datquality(data = sponges, dat.sim = sim.spo, Par = par.spo, transformation = "square root", method = "bray")

#Sampling and estimation of MultSE
samp.spo <- sampsd(sim.spo, par.spo, transformation = "square root",
                   method = "bray", n = 20, m = 20, k = 10)

#Summarizing results
sum.spo <- summary_ssp(results = samp.spo, multi.site = TRUE)

#Identification of optimal effort

opt.spo <- ioptimum(xx = sum.spo, multi.site = TRUE)

#plot
fig.2 <- plot_ssp(xx = sum.spo, opt = opt.spo, multi.site = TRUE)
fig.2

# Estimation of components of variation
dat<-sponges[,2:length(sponges)]

#Square root transformation of abundances
dat.t<-sqrt(dat)

#Bray-Curtys
library(vegan)
bc<-vegdist(dat.t, method = "bray")

#function to estimate components of variation in PERMANOVA
cv.permanova <- function(D, y) {
  D = as.matrix(D)
  N = dim(D)[1]
  g = length(levels(y[,1]))
  X = model.matrix(~y[,1])  #model matrix
  H = X %*% solve(t(X) %*% X) %*% t(X)  #Hat matrix
  I = diag(N)  #Identity matrix
  A = -0.5 * D^2
  G = A - apply(A, 1, mean) %o% rep(1, N) - rep(1, N) %o% apply(A, 2, mean) + mean(A)
  MS1 = sum(G * t(H))/(g - 1)  #Mean square of sites
  MS2 = sum(G * t(I - H))/(N - g)  #Mean square of residuals
  CV1 = (MS1 - MS2)/(N/g)# Components of variation of sites
  CV2 = MS2 # Components of variation of samples
  CV = c(CV1, CV2)
  sqrtCV = sqrt(CV)
  return(sqrtCV) #square root of components of variation
}

cv<-cv.permanova(D = bc, y = sponges)
cv
